# Emotional news affects social judgments independent of perceived media credibility

**DOI:** 10.1101/2020.02.29.971234

**Authors:** Julia Baum, Rasha Abdel Rahman

## Abstract

How does the credibility we attribute to media sources influence our opinions and judgments derived from news? Participants read headlines about the social behavior of depicted unfamiliar persons from websites of trusted or distrusted well-known German news media. As a consequence, persons paired with negative or positive headlines were judged more negative or positive than persons associated with neutral information independent of source credibility. Likewise, electrophysiological signatures of slow and controlled evaluative brain activity revealed a dominant influence of emotional headline contents regardless of credibility. Modulations of earlier brain responses associated with arousal and reflexive emotional processing show an effect of negative news and suggest that distrusted sources may even enhance the impact of negative headlines. These findings demonstrate that though we may have distinct perceptions about the credibility of media sources, information processing and social judgments rely on the emotional content of headlines, even when they stem from sources we distrust.

---

In times of massive online communication, news and information from various sources spreads rapidly, shaping personal opinions as well as public debates (Vosoughi et al., 2018). Aside from well-vetted news, intentionally or unintentionally spread misinformation, “fake news” and “alternative facts” have gained influence (Lazer et al., 2019). Despite the potentially detrimental effects of misinformation and their increasing prevalence in (social) media and political discourse, research on the consequences of being exposed to misinformation is scant, and little is known about the behavioral and neural correlates of processing information of questionable veracity (Baum et al., 2018). Experimental evidence revealing insights into the cognitive mechanisms can be vital to a comprehensive understanding of how we are affected by information from media (as argued, e.g., by Aral & Eckles, 2019; Lazer et al., 2018; and Vosoughi et al., 2018).

One resource-efficient and fast heuristic to assess the veracity of news is to consider the credibility of the source. Indeed, recent evidence suggests that we trust or distrust media sources based on criteria as familiarity, likability, social endorsement and reputation, and laypeople’s credibility assessments align with those of professional fact checkers (Metzger & Flanagin, 2013, Pennycook & Rand, 2018; Pennycook & Rand, 2019). However, despite our ability to evaluate the credibility of a source, little is known about the impact of such assessments on the cognitive processes underlying social judgments and decisions. The aim of the current study is to investigate the later (and possibly memory-related) consequences of having been exposed to news from various sources. Specifically, we asked how the perceived credibility of existing and well-known news sources affects subsequent information processing and social judgments based on person-related negative or positive headlines. We extracted event-related brain potentials (ERPs) from the electroencephalogram (EEG) to localize the effects and interactions of social-emotional information and source credibility at early reflexive and later more controlled processing stages to gain insight into the underlying cognitive mechanisms and brain signatures (please see Fig.1 for the study phases).

**Fig. 1:**
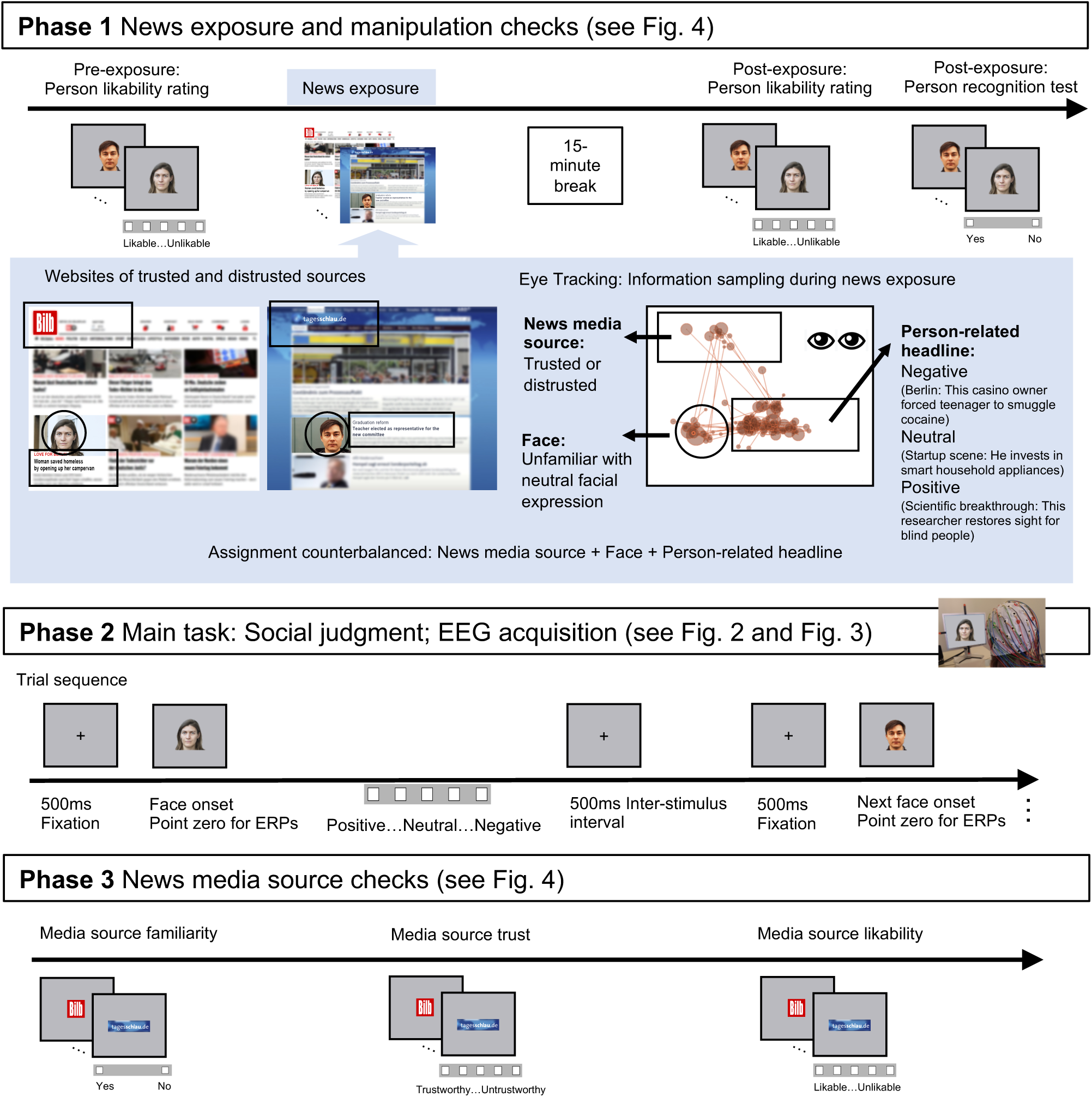
News exposure and manipulation checks before and after the main task.

When we are exposed to news we are confronted with verbal information (see Fig.1, Phase 1). Emotional person-related verbal information – even when minimal like in headlines – can change the affective value of people by mechanisms of verbal evaluative learning (also referred to as evaluative conditioning) as well as by attributional or propositional processes that may additionally take into account the relevance or truth-value of the information in its context (Bliss-Moreau et al., 2008; Ferrari et al., 2020; Mattarozzi et al., 2014; for a general review see De Houwer, Van Dessel, Moran; 2020). Some evidence of potential neural underpinnings of person-related verbal evaluative learning suggests that while emotional information may not affect very early visual processing (but see Galli et al., 2006), it can affect early and later conceptual processing that may rely on both, implicit and explicit memory of the information (Junghöfer et al., 2016; Kissler & Strehlow, 2012; and see introduction of ERP effects below). Yet, research on how these effects are modulated by the veracity of the information is scarce (Baum et al., 2018).

What are the expected consequences of having been exposed to emotional news from trusted and distrusted sources on social judgments (see Fig.1, Phase 2)? The family of dual-process theories distinguishes between two separate systems or interactive processes related to fast, impulsive, spontaneous and automatic processing on the one hand, and slower intentional and controlled processing on the other (e.g. Cunningham & Zelazo, 2007; Gawronski & Bodenhausen, 2006; Kahneman, 2003; Lieberman, 2007; Strack & Deutsch, 2004). This concept also relates to models of recognition and memory distinguishing faster and slower retrieval, with slower processes retrieving additional context and source information that may be stored unitized or separately (for a review see Yonelinas, 2002). For the memory-related processing in Phase 2, this suggests that our cognitive system initially spontaneously processes the emotional content of the headlines associated with the person irrespective of the credibility of the source, whereas later, more controlled processes should result in evaluations that take the credibility of the source into account, resulting in social judgments that are qualified according to the presumed credibility.

With respect to emotion processing, appraisal theories (Ellsworth & Scherer, 2003; Scherer, 2001) assume that stimuli are initially checked for a coarse detection of emotional salience, intrinsic pleasantness and arousal. This is followed by assessments regarding implications for the observer’s well-being, coping possibilities, and evaluations of the normative significance, like the compatibility with moral standards. This may also include the truth-value of information. Concerning the impact of news, and in analogy to dual process theories, emotional contents and source credibility should be processed at different points in time. While early emotional responses should be influenced only by the emotional content of headlines, later more controlled processes should take source credibility into account.

In ERPs fast and early processing has been related to an enhanced early posterior negativity (EPN) at about 200 – 300ms at occipito-temporal brain regions that indexes reflexive and arousal-related emotional processes (e.g., Junghöfer et al., 2001; Kissler et al., 2007; Schupp et al., 2003; Schupp et al., 2004). At later stages an enhanced late positive potential (LPP) at about 400 – 600ms at centro-parietal regions is associated with elaborate and reflective processing (Sabatinelli et al., 2013; Schacht & Sommer, 2009a; Schupp et al., 2004). Both components are sensitive to verbal affective person-related information associated with faces via verbal evaluative learning (for instance, EPN: Junghöfer et al., 2016; Luo et al., 2016; Suess et al., 2015; Wieser et al., 2014; Xu et al., 2016; LPP: Abdel Rahman, 2011; Baum et al., 2018; Luo et al., 2016; Kissler & Strehlow, 2012). Crucially, the LPP is sensitive to additional information such as context and relevance, putting emotional contents into perspective (Blechert et al., 2012; Herbert et al., 2011; 2013; Rellecke et al., 2012; Schacht & Sommer, 2009b; Schindler et al., 2019), whereas the EPN is relatively independent of task demands and the relevance of emotional contents in a given context (Herbert et al., 2011; Herbert et al., 2013; Schacht & Sommer, 2009b). It is noteworthy that this evidence of additional contextual influences on ERPs comes from studies testing effects of emotional information immediately, while there is scarce evidence of such contextual effects on later consequences (Baum et al., 2018). We expected that the EPN is mainly sensitive to the emotional content of the headlines irrespective of source credibility, whereas emotion effects in LPP amplitudes should be modulated by source credibility, with reduced amplitudes for distrusted sources.

To summarize, based on dual-process theories distinguishing fast impulsive and slower more controlled processes, we expected that early processing of faces associated with emotional vs. neutral headlines from trusted and distrusted sources should be modulated only by effects of emotion, whereas later controlled evaluation should take source credibility into account, resulting in tempered social judgments. This modulation may be primarily found for positive headlines if negative information is prioritized as protection against potential threat (cf. Baum et al., 2018). The present study was preregistered under the OSF (Baum & Abdel Rahman, 2018^2^).

Fig. 1. Overview of the well-controlled experimental study design with three phases. In **Phase 1** participants were exposed to experimental but authentic websites of existing and widely distributed mainstream German news media (e.g. *Tagesschau* or *Bild*) that were selected based on their pre-rated high or poor credibility. English-speaking analogies may be the *BBC, The Guardian*, or *The New York Times* on the one hand and *Fox News, Daily Mirror*, or *The Sun* on the other. Each website presented the news media source logo, the face, and the headline containing negative, positive, or neutral emotional person-related information; all other details were blurred (in the experiment original layouts, logos and fonts were used). To enhance authenticity we added news reports about well-known persons as fillers. The assignment of unfamiliar faces to conditions was counterbalanced: while one participant was exposed to each face only in one context condition, the faces were presented equally often in each condition across participants. An additional eye tracking experiment with different participants verified the sampling of source information during news exposure (shown here: example data of one participant for one website, lines represent saccades, points represent fixations and point magnitude their duration). To check whether the news exposure manipulation was successful, we subsequently tested whether the faces were reliably recognized and how likable participants found each person before and after news exposure. In **Phase 2**, the main experimental task followed, in which the faces were presented in isolation and the EEG was registered while participants judged the depicted persons based on the information they had been exposed to (social judgment). Just as it is typically the case when reading news headlines, participants were not explicitly instructed to consider the credibility of the source. Instead, they were asked to make their judgment based on the information from Phase 1. In **Phase 3** (after the main task), participants rated the familiarity, likability and credibility of the news media sources as an additional manipulation check.

## Method

### Participants

The sample size was preregistered and planned according to the requirements of the counterbalancing and based on power analyses, see SI-page 1. The final data set consisted of thirty participants (*M*_age_=25 (*SD*=5.36), 25 females, all right-handed). Four participants were excluded (one was familiar with face databases, two rated the trustworthiness equal across sources, one did not acquire person-related information) and replaced with new participants. Participants were compensated in form of course credit or money. They were (de)briefed about the procedures and signed informed consent. The study was approved by the local ethics committee.

### Materials

Websites of news media combined source, face, and headline (for examples see Fig.1, Phase 1). We edited each colored face photograph onto a natural background (e.g., street scene, wall), inserted it onto the website and changed the headline via source code, keeping the characteristic font (with font size kept similar across media sources). Thus we maintained the distinctive layout of the media sources while experimentally manipulating the content, since the layout and visual design of websites plays an important role in assessing the credibility of a source (Metzger et al., 2013). In Phase 1, website screen shots were displayed full screen and showed the prominent logo on the top of the page, the face, and the headline, while all other details were blurred. For the news exposure, 24 unfamiliar faces were equally assigned to neutral, negative and positive headlines, with counterbalanced assignment across participants. The assignment of faces and headlines to media sources was also counterbalanced across participants, with 12 target faces appearing in trusted sources and 12 faces in distrusted sources, resulting in 4 target faces in each condition of the 3×2 design. Affective information for 8 well-known filler faces referred to recent news about them and the assignment of headlines was fixed for all participants.

News media sources were selected based on pre-ratings of credibility and familiarity with a different group of German participants (N=38, 33 females, *M*_age_=26 (*SD*=4.69), all students). The pre-rating tested 35 German news media sources, including well-known, less-well-known, and highly-partisan sources. The rating scale was from 3 (very credible) to -3 (not credible). We selected the four sources rated as most credible (*M*=1.77, 95%-CI [1.57, 1.97]), and the four rated as least credible (*M*=-1.64, 95%-CI [-1.92, -1.37]), all highly familiar (familiar=1, unfamiliar=0; *M*=.98 for trusted and distrusted). Credibility ratings were significantly higher for trusted than for distrusted sources, *t*(37)=14.83, *p*<.001. Colored screen shots of the sources’ logos were presented in similar size in the media source ratings of the current experiment (2.7×3.5cm).

Face stimuli were colored frontal portraits of 24 unfamiliar faces with neutral facial expressions, presented on a grey background during the main task and manipulation checks (2.7×3.5cm, viewing distance 70cm; from multiple databases, see SI-page 14). Eight familiar filler faces (e.g. Emma Watson, Harvey Weinstein) were added to make the target persons’ existence credible.

Headlines describing social behavior were either neutral, negative, or positive (for all headlines see SI-Table S20). Pre-ratings with different participants confirmed their valence and showed that positive and negative headlines were equally more arousing than neutral headlines (see SI-page 14).

### Procedure

The procedure entails three phases (Fig.1) as a variant of a well-established design (cf. Abdel Rahman, 2011; Baum et al., 2018; Suess et al., 2015). In Phase 1, the experiment started with a person likability rating of all faces on a 5-point scale (pre-exposure rating). Response buttons were placed in front of participants. Then the news exposure followed. Participants were instructed as follows: “You now receive information of various kinds about these people, taken from media reports. Unrelated content and details remain unrecognizable. Please read the information carefully”. Each trial started showing the website –which was blurred except for the logo of the media source– for one second. For the remaining 5s, the logo, the face and the headline were unblurred. Websites were presented in blocks of 8, including all experimental conditions and 2 fillers. Each website was presented 5 times in total (160 trials in total). To keep participants engaged with the task, they occasionally answered short yes-or-no questions about the persons, e.g. *Is the behavior of this person common?* (asked in about 22% of the trials of Phase 1). After completion of the news exposure, participants had a 15-minute break. Phase 1 ended with a post-exposure likability rating (see earlier) and a recognition test as manipulation checks. In the recognition test participants decided whether a face had been encountered in the news exposure or not (this included 32 additional unfamiliar filler faces).

In Phase 2 the EEG was recorded while a social judgment task was employed as the main task. Participants judged how negative, neutral, or positive the depicted person was based on information acquired in Phase 1. Participants judged on a 5-point scale, enabling them to nuance their answers between neutral and negative / positive. To enhance the signal-to-noise ratio necessary for the EEG data quality, the task was repeated 20 times block-wise, separated by breaks, resulting in 80 trials per condition (excluding fillers). Participants were told that the repetition of the task is a technical necessity for EEG measurements. Trials started with a 500ms pre-stimulus fixation cross and had a 500ms inter-trial-interval. Faces were presented until response or for a maximum of 3s.

Phase 3 entailed manipulation checks of the media sources. First, participants saw the logos and were asked if they knew the sources. Then they rated how trustworthy they consider each source, on a 5-point scale from trustworthy to untrustworthy while the EEG was recorded. The trust rating was repeated 10 times, resulting in 40 trials per condition and logos were presented until response. At last, participants were asked to rate how likeable they find each media source. This rating was included because likability may not necessarily be equivalent with credibility (e.g. one may enjoy reading gossip papers, without trusting its contents).

The direction of scales was counterbalanced, i.e. there were two versions, in version one the 5 buttons ranged from positive (left) to negative (right), and in version two from negative (left) to positive (right). This was consistent for all tasks and phases, i.e. very likeable, positive, yes, and very credible on the left for version one and vice versa for version two. After the experiment, participants were asked to reproduce the contents of the headline about each person to check if they remembered the broad information. Phase 1 lasted 30minutes, Phase 2 and 3 together 40minutes and participants were compensated for all time spent at the lab.

### EEG Data recording and preprocessing

The EEG was recorded with BrainAmpDC amplifiers, from 62 Ag/AgCl-electrodes as specified by the extended 10-20 system, referenced to the left mastoid with FCz as Ground Electrode. Impedance was kept under 5kΩ. EEG data was recorded at a sampling rate of 5kHz and down-sampled to 500Hz using a low-cutoff of 0.016Hz and a high-cutoff of 1000Hz. Horizontal and vertical electrooculograms were obtained with peripheral electrodes at the left and right canthi of both eyes, and above and below the left eye. A short calibration procedure traced individual eye movements after the experiment, that were later used to correct for eye movement artifacts.

Offline, the continuous EEG was transformed to average reference and low-pass filtered at 30Hz pass-band edge (zero-phase FIR-filter with transition band width of 7.5Hz and cutoff frequency (−6dB):33.75Hz, EEGlab-toolbox version 13_5_4b; Delorme & Makeig, 2004). Using BESA (Berg & Scherg, 1991), we removed artifacts due to eye movements by applying a spatiotemporal dipole modeling procedure for each participant individually. Trials with remaining artifacts were rejected, i.e. trials with amplitudes over ±200µV, changing more than 50µV between samples or more than 200µV within single epochs, or containing baseline drifts. Error- and artifact-free EEG data was segmented into epochs of 1s, starting 100ms prior to stimulus onset, with a 100ms pre-stimulus baseline. For EEG analysis, per participant an average of 79 trials per condition remained (range: 73-80) and in each condition 98% of trials were kept overall (neutral-trusted 2357, neutral-distrusted 2364, negative-trusted 2350, negative-distrusted 2355, positive-trusted 2363, positive-distrusted 2362). Trials where no judgment was given were excluded (in the social judgment task that were 33 out of 14400).

### Data analysis

ERP analyses focus on two regions of interest (ROI), the EPN (at electrode sites PO7, PO8, PO9, PO10, TP9, TP10, 200–350ms after face stimulus onset) and the LPP component (Pz, CPz, POz, P3, P4, 400–600ms), based on previous findings of emotional stimulus content (e.g. Schupp et al., 2003) and affective information (e.g. Abdel Rahman, 2011; Baum et al., 2018). To explore effects occurring during early visual face processing, we additionally analyzed the P100 (PO3, PO4, O1, O2, 100–150ms), and the N170 (P7, P8, PO7, PO8, 150– 200ms), based on previous findings (e.g. Abdel Rahman & Sommer, 2012). P100 and N170 results are available in the SI-Tables S7-S9. Amplitudes were averaged over ROIs and time windows on single-trial level.

We used mixed-effects regression models on single-trial data of behavioral measures and ERPs (Frömer et al., 2018). For continuous dependent variables we used linear mixed models (LMMs; Bates et al., 2015b: *lme4* v.1.1-17 in R) and tested the significance of fixed effects coefficients (p-value < .05) by Satterthwaite approximation (*summary* function of *lmerTest* v.3.0-1, Kuznetsova et al., 2017). For ordinal dependent variables we used cumulative-link mixed-models fitted with Laplace approximation (CLMMs; *ordinal* v.2019.12-10, Christensen, 2019)^3^. For each dependent variable, the model was specified with fixed effects for the experimental factors *headline content* (negative, positive, neutral; with neutral as the reference level) and *source credibility* (trusted, distrusted; with distrusted as the reference level) and their interaction. Both factors were modeled as repeated contrasts that compare the means of factor levels to the respective reference level. Thus coefficients represent our hypotheses that expect emotion effects of negative vs. neutral and of positive vs. neutral headline content, each in interaction with source credibility, with reduced or absent effects of headline content for distrusted sources (see Schad et al., 2020 for details on testing a-priori hypotheses through contrast specification in LMMs). We fitted models with a maximal crossed random-effects structure correcting for by-subjects and by-face-stimuli random intercepts and slopes. If necessary, random-slopes correlation parameters were set to zero and slopes explaining zero variance were omitted to achieve convergence and avoid overparameterization (Bates et al., 2015a; final random structures are reported in the results Tables). To test our hypotheses that emotion effects may be present only for trusted but not distrusted sources, we tested emotion effects separately for each source credibility condition as a follow-up (via *emmeans* v.1.4.6, Lenth, 2020, with false-discovery-rate adjusted p-values, Benjamin & Hochberg, 1995; see Tables 2 and 4). We report point estimates (*b*), 95% confidence intervals for LMMs, standard errors, t-values for LMMs, z-values for CLMMs, and p-values for the fixed effects coefficients. Data and code can be accessed online^4^.

## Results

### Effects of Emotional News on Information Processing and Social Judgments (Phase 2)

#### Behavioral results

Persons associated to negative headlines were judged as more negative relative to persons associated to neutral headlines, and judgments based on negative headlines were faster than when based on neutral headlines (please see Table 1 and Fig.2). Source credibility did not modulate the negative headline effects in judgment decisions and latencies (Table 1). Unexpectedly, social judgments based on negative vs. neutral headlines were more negative and faster for both, trusted and distrusted sources (Table 2).

**Fig. 2:**
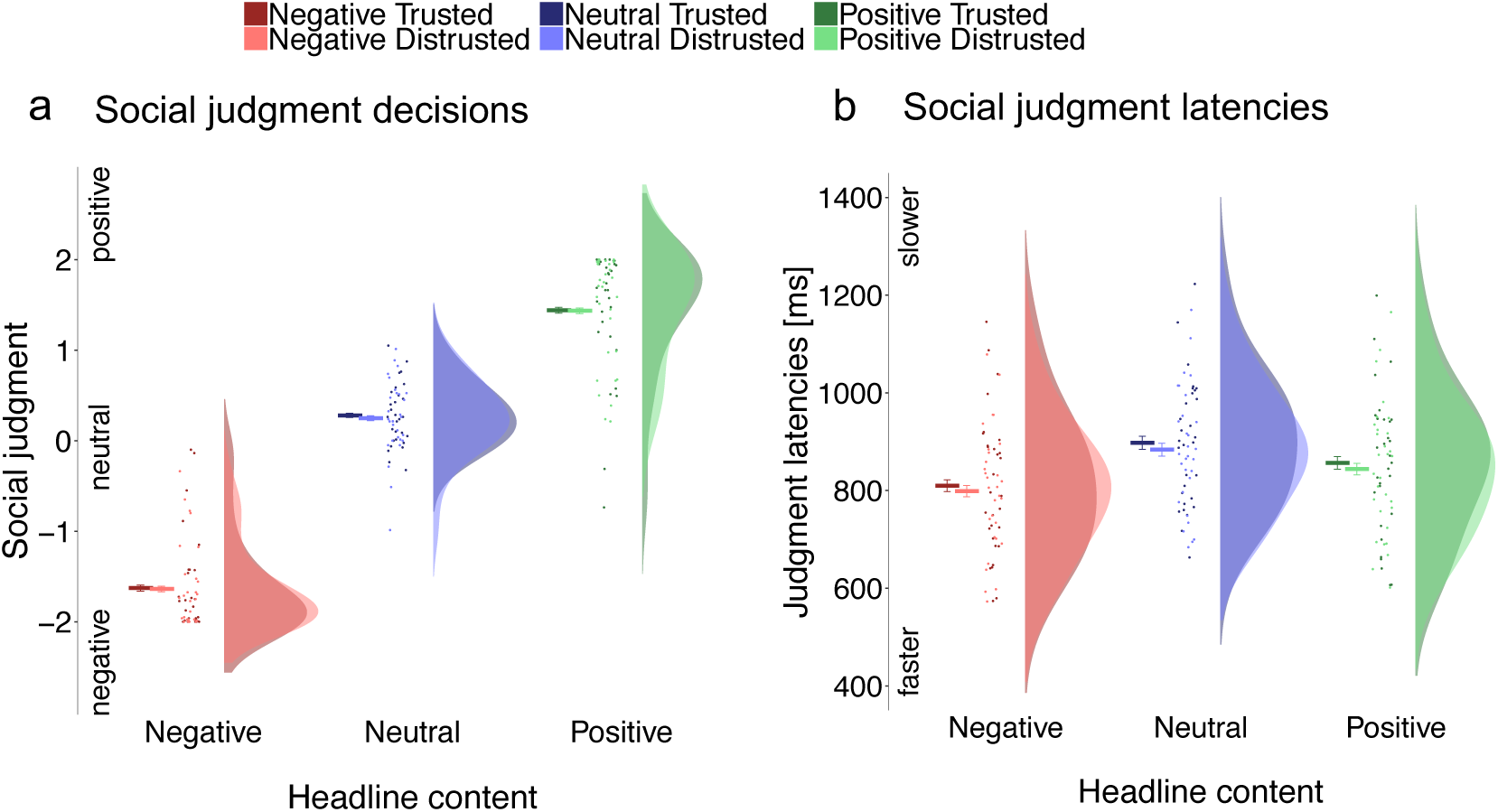
**Phase 2** Main task: Social judgment behavioral results.

**Table 1.** Mixed model summary statistics show effects of source credibility, negative and positive headline content and their interactions on behavioral dependent variables in the social judgment task. Effects on social judgment decisions and latencies were estimated in separate mixed models and fixed effects were coded as repeated contrasts according to our hypotheses.

| *Coefficient* | **Social Judgment Decisions** |  |  |  | **Social Judgment Latencies**<br>**[-1000/latency(ms)]** |  |  |  |
| --- | --- | --- | --- | --- | --- | --- | --- | --- |
|  | *b* | *SE* | *z* | *p* | *b* (95%-CI) | *SE* | *t* | *p* |
| Intercept (Grand Mean) |  |  |  |  | -1.31 (-1.37 – -1.25) | 0.03 | -44.45 | <0.001 |
| Source Credibility (Trusted vs. Distrusted) | -0.01 | 0.39 | -0.02 | 0.984 | 0.01 (-0.01 – 0.04) | 0.01 | 0.84 | 0.410 |
| Negative Headline Content (Neg vs. Neu) | -8.20 | 0.75 | -11.00 | <0.001 | -0.13 (-0.18 – -0.07) | 0.03 | -4.69 | <0.001 |
| Source Credibility * Negative Headline Content | -0.28 | 1.11 | -0.26 | 0.799 | -0.02 (-0.08 – 0.04) | 0.03 | -0.65 | 0.521 |
| Positive Headline Content (Pos vs. Neu) | 4.93 | 0.61 | 8.07 | <0.001 | -0.06 (-0.10 – -0.02) | 0.02 | -3.04 | 0.004 |
| Source Credibility * Positive Headline Content | -0.05 | 0.46 | -0.12 | 0.908 | -0.02 (-0.08 – 0.05) | 0.03 | -0.55 | 0.583 |
| Model Formula | Decision ~ Headline Content * Source Credibility<br>+ (S+Neg+S*Neg+Pos+S*Pos \|\| subject)<br>+ (S+Neg+S*Neg+Pos+S*Pos \|\| face) |  |  |  | Latency ~ Headline Content * Source Credibility<br>+ (S+Neg+S*Neg+Pos+S*Pos \|\| subject)<br>+ (S+Neg+S*Neg+Pos+S*Pos \|\| face) |  |  |  |

*Note.* Double bars in random effects terms represent correlation parameters set to zero. Abbreviations for slopes in the random effects terms: S=Source Credibility, Neg=Negative Headline Content, Pos=Positive Headline Content. Face stands for face stimulus.
CLMM threshold coefficients for social judgment decisions: -2 | -1: $b=-3.9$ , $SE=.07$ , $z=-57.9$ ; -1 | 0: $b=-1.75$ , $SE=.05$ , $z=-33.2$ ; 0 | 1 $b=2.0$ , $SE=.05$ , $z=40.4$ ; 1 | 2: $b=5.1$ , $SE=.07$ , $z=74.3$

**Table 2.** Negative and positive headline content effects on social judgment decisions and latencies separately within each source credibility condition computed from the models in Table 1.

| *Contrast* | **Social Judgment Decisions** |  |  |  | **Social Judgment Latencies**<br>**[-1000/latency(ms)]** |  |  |  |
| --- | --- | --- | --- | --- | --- | --- | --- | --- |
|  | *b* | *SE* | *z* | *p* | *b* | *SE* | *t* | *p* |
| Trusted: Neg vs. Neu | -8.35 | 0.93 | -8.98 | <0.001 | -0.14 | 0.03 | -4.39 | <0.001 |
| Distrusted: Neg vs. Neu | -8.06 | 0.93 | -8.67 | <0.001 | -0.12 | 0.03 | -3.74 | <0.001 |
| Trusted: Pos vs. Neu | 4.91 | 0.65 | 7.50 | <0.001 | -0.07 | 0.03 | -2.71 | 0.011 |
| Distrusted: Pos vs. Neu | 4.96 | 0.65 | 7.60 | <0.001 | -0.05 | 0.03 | -2.00 | 0.049 |

For positive headlines, social judgments were more positive and also faster compared to neutral headlines (Table 1). These effects were not modulated by source credibility (Table 1). Social judgments of positive vs. neutral headlines were more positive and faster for trusted and distrusted sources (Table 2).

Post-hoc (non-preregistered), we included repetition as a covariate to test whether social judgments and their latencies were biased towards focusing on emotional contents by repeating the task, which was necessary to ensure EEG data quality. The three-way interactions were not significant (all *ts* <|.9|, all *ps*>.4; see SI-Table S3). Moreover, testing only the first judgments per face (task was repeated block wise) resulted in the same pattern (SI-Table S4). We conclude that repetition did not change the result pattern.

Fig. 2. In **Phase 2** the social judgment was performed as main task to investigate the effects of emotional news and source credibility. Behavioral results show that **a)** persons were judged based on emotional headline content, whereas source credibility had no influence. **b)** Judgments based on emotional headlines were faster than neutral, but not tempered by source credibility. Raincloud plots (Allen et al., 2019) show means and 95% confidence intervals calculated with the *summarySEwithin* function (Morey, 2008) on single trial data, with points and distributions for data aggregated by subject.

#### Event-related brain potentials

##### EPN

To investigate relatively fast and reflexive emotional processing we focused on the EPN component. Negative compared to neutral headlines elicited an enhanced negativity, and there was a trend for an interaction with source credibility (please see Table 3). The EPN effect of negative headlines was enhanced for distrusted sources, but absent for trusted sources (Table 4 and Fig.3a,c).

**Table 3.** LMM summary statistics show effects of source credibility, negative and positive headline content and their interactions on ERPs as dependent variables in the social judgment task. Effects on the predefined ROI and time range of the EPN and LPP amplitudes were estimated in separate LMMs and fixed effects were coded as repeated contrasts according to our hypotheses.

| *Coefficient* | **EPN** |  |  |  | **LPP** |  |  |  |
| --- | --- | --- | --- | --- | --- | --- | --- | --- |
|  | *b* (95%-CI) | *SE* | *t* | *p* | *b* (95%-CI) | *SE* | *t* | *p* |
| Intercept (Grand Mean) | 2.41<br>(1.33 – 3.49) | 0.55 | 4.36 | <0.001 | 4.64<br>(3.88 – 5.41) | 0.39 | 11.85 | <0.001 |
| Source Credibility (Trusted vs. Distrusted) | -0.02<br>(-0.23 – 0.18) | 0.11 | -0.23 | 0.819 | 0.10<br>(-0.11 – 0.31) | 0.11 | 0.95 | 0.353 |
| Negative Headline Content (Neg vs. Neu) | -0.29<br>(-0.50 – -0.08) | 0.11 | -2.65 | 0.014 | 1.13<br>(0.80 – 1.45) | 0.17 | 6.79 | <0.001 |
| Source Credibility * Negative Headline Content | 0.42<br>(0.01 – 0.84) | 0.21 | 2.00 | 0.056 | 0.36<br>(-0.06 – 0.78) | 0.21 | 1.69 | 0.101 |
| Positive Headline Content (Pos vs. Neu) | -0.11<br>(-0.32 – 0.09) | 0.10 | -1.09 | 0.287 | 0.50<br>(0.23 – 0.77) | 0.14 | 3.60 | 0.001 |
| Source Credibility * Positive Headline Content | 0.14<br>(-0.33 – 0.61) | 0.24 | 0.59 | 0.559 | 0.21<br>(-0.29 – 0.71) | 0.26 | 0.83 | 0.414 |
| Model Formula | EPN ~ Headline Content * Source Credibility<br>+ (S+Neg+S*Neg+Pos+S*Pos \|\| subject)<br>+ (S+Neg+S*Neg+Pos+S*Pos \|\| face) |  |  |  | LPP ~ Headline Content * Source Credibility<br>+ (S+Neg+S*Neg+Pos+S*Pos \|\| subject)<br>+ (S+Neg+Pos+S*Pos \|\| face) |  |  |  |

*Note.* Double bars in random effects terms set correlation parameters to zero. Abbreviations for slopes in the random effects terms: S=Source Credibility, Neg=Negative Headline Content, Pos=Positive Headline Content. Face stands for face stimulus.

**Table 4.** Negative and positive headline content effects on EPN and LPP separately within each source credibility condition computed from the models in Table 3.

| *Contrast* | **EPN** |  |  |  | **LPP** |  |  |  |
| --- | --- | --- | --- | --- | --- | --- | --- | --- |
|  | *b* | *SE* | *t* | *p* | *b* | *SE* | *t* | *p* |
| Trusted: Neg vs. Neu | -0.08 | 0.15 | -0.52 | 0.79 | 1.31 | 0.20 | 6.62 | <0.001 |
| Distrusted: Neg vs. Neu | -0.50 | 0.15 | -3.30 | 0.007 | 0.95 | 0.20 | 4.79 | <0.001 |
| Trusted: Pos vs. Neu | -0.04 | 0.16 | -0.27 | 0.786 | 0.60 | 0.19 | 3.20 | 0.031 |
| Distrusted: Pos vs. Neu | -0.18 | 0.16 | -1.16 | 0.501 | 0.39 | 0.19 | 2.07 | 0.043 |

**Fig. 3:**
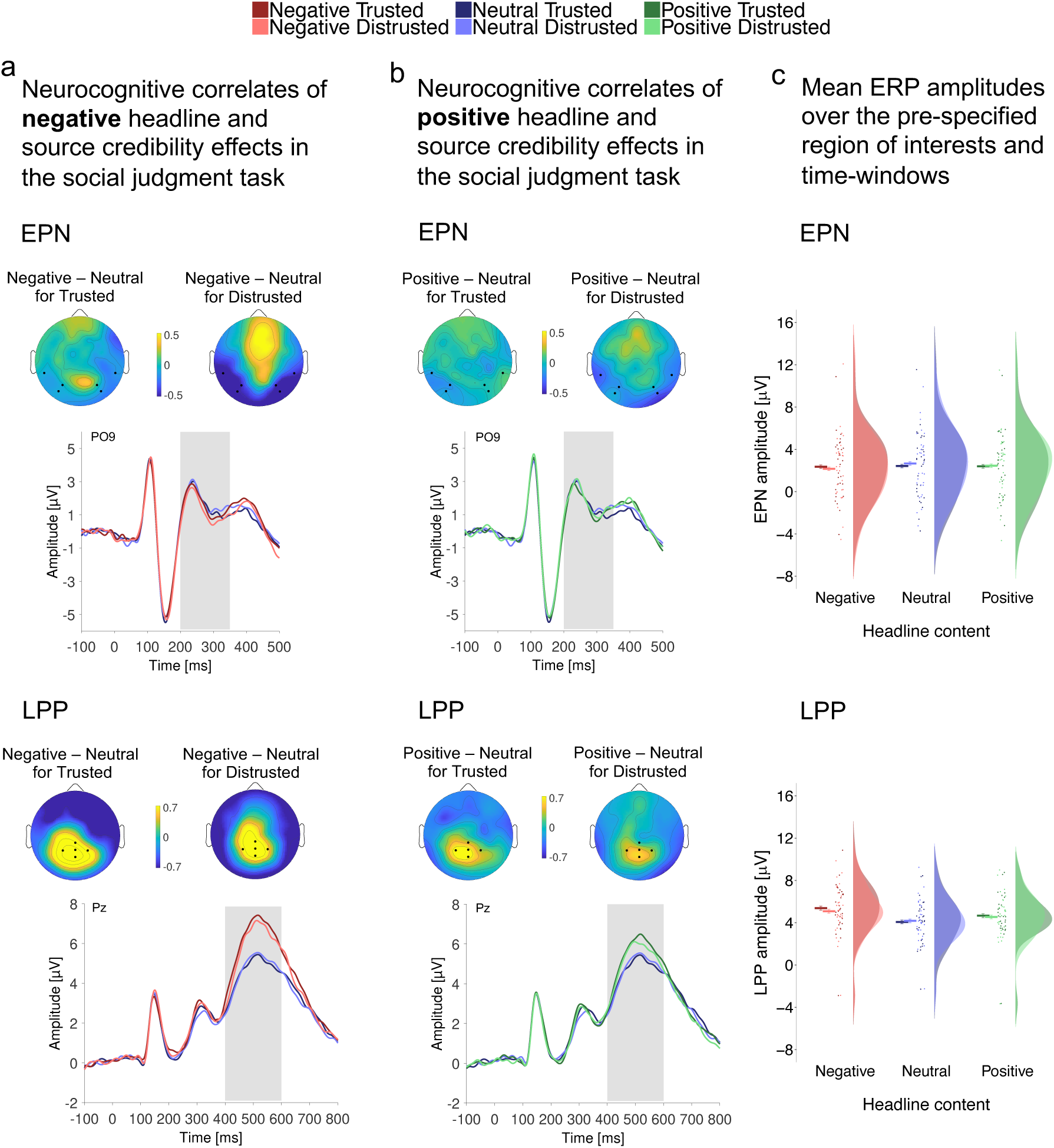
**Phase 2** Main task: Social judgment EEG results.

For positive headlines, we found no EPN effect for positive compared to neutral headlines, no interaction with source credibility (Table 3), and no EPN effects nested in trusted or distrusted sources (Table 4 and Fig. 3b,c).

##### LPP

To investigate more controlled evaluative processing, we tested effects in the later LPP component. For negative headlines, we found an enhanced LPP compared to neutral headlines and no interaction with source credibility (Table 3). Negative information from both, trusted and distrusted sources elicited LPP effects (Table 4 and Fig.3a,c).

For positive headlines, the LPP was enhanced compared to neutral headlines and there was no interaction with source credibility (Table 3). Positive information from trusted and distrusted sources elicited LPP effects (Table 4 and Fig.3b,c).

Post-hoc (non-preregistered), we included judgment latencies as a covariate to account for motor responses in the LPP results. This did not change the effects of predictors and three-way-interactions were not significant (all *ts* <1, all *ps*>.3; see SI-Table S6). We cannot fully exclude the possibility of motor-response or -preparation influences. Yet, we consider motor response confounds unlikely because first, all trials involved motor responses (Luck, 2014) and second, latency differences were taken into account in the model. Thus, mostly unsystematic or non-linear motor response-related differences could have affected the LPP.

Fig. 3. In **Phase 2** the EEG was acquired while social judgments were performed to investigate the neurocognitive correlates of emotional news and source credibility effects. **a)** ERP results for persons related to negative headline content reveal that reflexive emotional processing in the EPN (200–350ms) was affected by headline content. Evaluative processing in the LPP (400–600ms) was enhanced for negative headlines from trusted as well as distrusted sources. **b)** For persons related to positive headlines no EPN (200–350ms) modulation was observed, and the LPP (400–600ms) was enhanced for positive headlines from trusted and distrusted sources. In **a**, **b**, grand average ERPs are shown for the EPN at electrode sites PO9 and for the LPP at Pz, and scalp distributions show the effects as differences between conditions in the respective time windows shaded in grey. **c**) Mean ERP amplitude sizes are shown for the pre-specified regions-of-interest and time window of the EPN and LPP. Raincloud plots (Allen et al., 2019) show means and 95% confidence intervals calculated with the *summarySEwithin* function (Morey, 2008) on single trial data, and points, boxplots, and distributions for data aggregated by subject.

### News Exposure and Manipulation Checks (Phase 1)

We manipulated headline content and news media credibility during news exposure and demonstrate that these manipulations were successful (see Fig.4a,b). Pre-exposure person likability ratings were on average neutral (SI-Tables S10-S12), whereas after exposure persons were disliked when associated with negative headlines and liked when associated with positive headlines (b=-1.52, 95%-CI [-1.73, -1.31], *t*=-13.96, *p*<.001 and b=.78, 95%-CI [.61, .95], *t*=9.01, *p*<.001, respectively). Source credibility did not modulate likability ratings (*ts* <|.97|, *ps*>.3). In the post-exposure recognition test, faces were successfully recognized across conditions, *M*=97.3%. There were no effects of headline or source on accuracy (SI-Tables S13,S14).

**Fig. 4:**
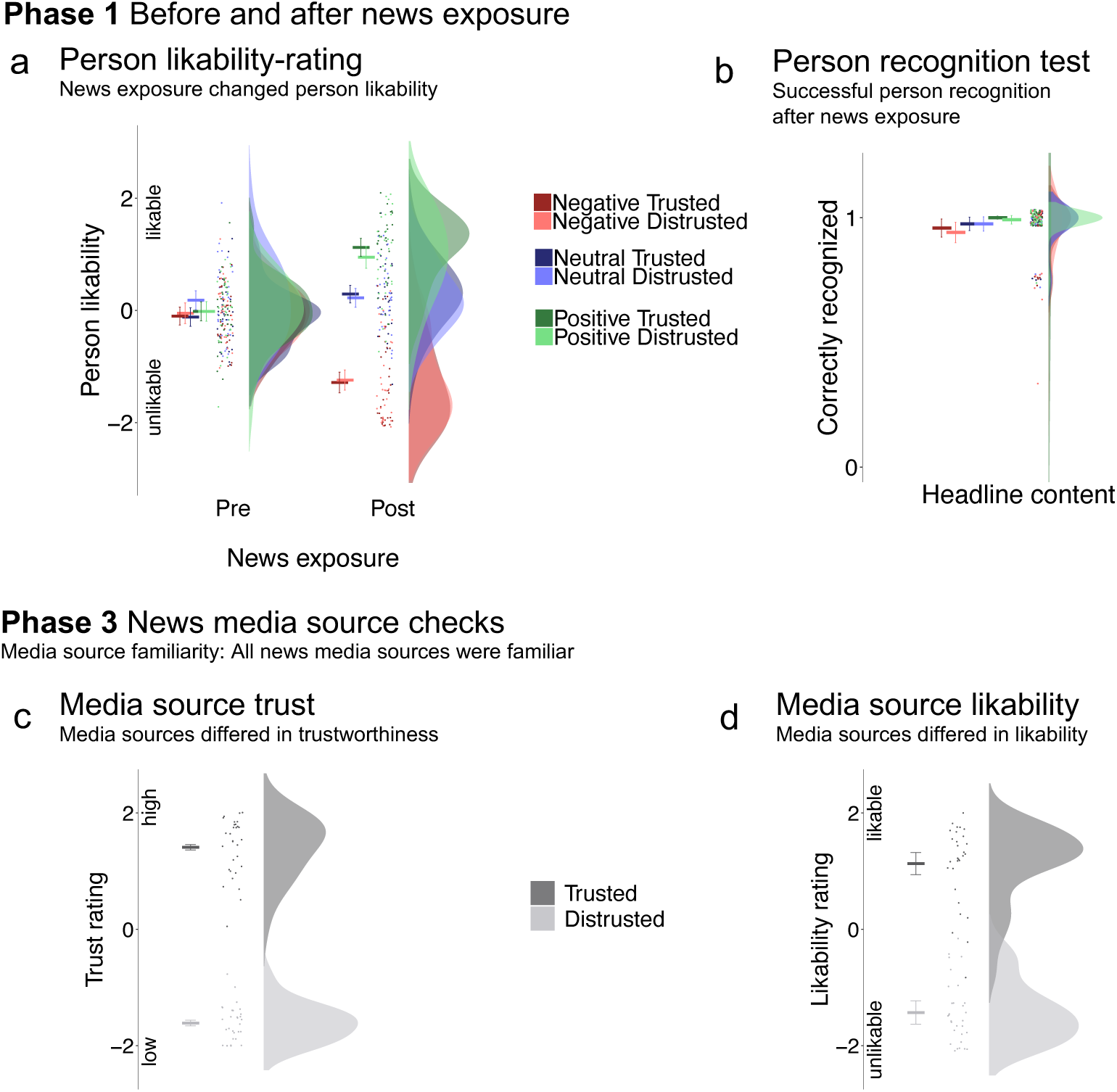
Results of manipulation checks before and after the main task.

We conducted an additional eye-tracking experiment with different participants (N=12, M_age_=25 (*SD*=7.93), 8 females) to check whether participants acknowledge the media source during news exposure, without having been explicitly instructed (see Procedure). One-sample-t-tests confirmed that per face presented in the website context the source fixation durations and frequencies on the source logo were above zero (*M*=896ms, 95%-CI [440,-]; *t*(11)=3.53, *p*=.002, d=1.02 and *M*=4.1, 95%-CI [2.2,-]; *t*(11)=3.93, *p*=.001, d=1.14; see Fig.1 and SI-page 11). Furthermore, we tested if the blurred layout in itself provides cues of the media source. In a separate task after news exposure participants assigned screenshots of websites where the logo had been removed to one of two sources (correct media source vs. logo of a different source from the other credibility condition). 90% of the layouts were correctly identified (*M*=.90, 95%-CI [.86,-], *t*(11)=40.58, *p*<.001, d=11.71).

### News Media Source Checks (Phase 3)

All participants were familiar with all media sources. Distrusted sources were rated as untrustworthy and less likable, whereas trusted sources were rated as trustworthy and likable (source credibility effect in trust ratings: b=3.02, 95%-CI [2.66, 3.38], *t*=16.64, *p*<.001, in likability ratings: b=2.56, 95%-CI [2.09, 3.02], *t*=10.80, *p*<.001; see Fig.4c,d and SI-page 11f).

Fig. 4. In **Phase 1** pre- and post-exposure person likability ratings and a post-exposure person recognition test served as manipulation checks for the news exposure. **a)** Persons were liked or disliked depending on the associated headline content, unaffected by source credibility. **b)** Persons were successfully recognized equally across conditions. In **Phase 3** news media source checks confirmed that all sources were familiar, and that they were differentiated in **c)** trustworthiness and **d)** likability. Raincloud plots (Allen et al., 2019) show means and 95% confidence intervals calculated with the *summarySEwithin* function (Morey, 2008) on single trial data, with points and distributions for data aggregated by subject.

## Discussion

Here we show that emotional person-related news headlines strongly affect subsequent information processing and social judgments irrespective of whether the source is perceived as credible or not. Emotional contents of headlines determined social judgments and affected slow evaluative brain responses in the LPP component known to be sensitive to context information and deliberate control. Crucially, none of these effects was modulated by source credibility, suggesting that headlines in news media may have an even stronger than expected influence on information processing and social judgments. Indeed, even if we assume that there are subtle traces of source credibility modulations that are difficult to detect, the fact remains that headlines from distrusted sources induce strong and robust effects of emotional information on social judgments.

Fast emotional brain modulations in the EPN component associated with arousal and sensation-related reflexive processing were modulated by emotional headline content and show furthermore that, if anything, distrusted sources may even enhance, instead of reduce, the impact of negative compared to neutral headlines. Please note however that this early interaction of headline content and source credibility was not predicted and the interaction was only marginally significant, even though clear and robust emotion effects were found only for distrusted sources. Future evidence should reveal additional evidence on the scope and limits of this effect. We speculate that this influence specifically of negative (but not positive) social-emotional information from distrusted sources may explain in part the popularity and success of (media) sources of questionable credibility: Untrustworthy negative social information may induce even positive states of enhanced arousal or excitation (cf. Menninghaus et al., 2017), increasing the impact of negative information (cf. Kahneman & Tversky, 1979; Zillmann, 2008). Indirect evidence for a possible compounded effect of source and headline comes from research demonstrating that arousal induced by irrelevant contexts (e.g. vocal affect) can change the subsequent emotional evaluation of neutral words, an instance of evaluative conditioning (Schirmer, 2010). Taken together, we conclude that low levels of perceived credibility may, if anything, even enhance the early reception of negative headlines. As discussed above, this may be due to pleasant states of arousal associated with untrustworthy negative information (gossip) or due to a form of evaluative learning resulting in negative affect.

The trend for an EPN modulation is unlikely to be affected by the differences in perceptual salience of the different source conditions because the faces were presented in isolation during social judgment.

The present effects were observed even though participants clearly distinguished between trusted and distrusted sources, as reflected in different measures. First, the perceived credibility of the news sources was determined in a separate rating study, which was confirmed by the participants of the present study, and early emotional responses in the EPN were induced by the logos of media sources judged as untrustworthy relative to trustworthy sources (Phase 3, see SI-page 13). Please note the EPN elicited by logos is likely biased by the real-life differences in perceptual salience (e.g. red vs. blue). Third, active eye movements in an additional manipulation check study demonstrate that the media sources of the headlines are actively acknowledged during news exposure. Finally, we found that even the blurred website layouts without logos provide reliable cues of the source and its credibility. We are therefore confident that the credibility of media sources was successfully manipulated and noticed by the participants.

The pattern of results is in contrast to our theoretical predictions, assuming that fast reflexive processes are mainly based on the emotional contents of the headlines, whereas slower, more controlled evaluations reflected in the LPP component and the actual judgments are modulated by source credibility, putting emotional information of questionable credibility into perspective. In contrast, our findings are in line with recent evidence of strong emotion effects of untrustworthy affective person-related information. In a related study we manipulated the trustworthiness of person-related information with verbal markers such as *supposedly, people assume* etc. (e.g. *He allegedly bullied his trainee*; Baum et al., 2018). Verbal qualifiers have an important communicative and legal function to indicate that the information might not be truthful. Just like in the present study, while participants understood the questionable veracity, person judgments and evaluative brain responses were determined by the emotional information independent of the verbally marked trustworthiness. The similarity of the findings may suggest a general mechanism.

The use of a controlled experimental design with a systematic manipulation of source credibility offers full control of confounding factors such as visual differences between faces, but it also differs in many ways from natural situations. However, here we presented existing and well-known media sources that are stored in long-term memory, including their perceived credibility. This should have even strengthened credibility effects. As in real-life situations when confronted with emotional headlines containing social information, participants in our experiment were not instructed to actively suppress the emotional content or to contemplate about the credibility of the source, but were free to consider source credibility to put their judgments into perspective. In the main task, we asked participants to repeatedly judge the person, which may have induced a strong focus on the news contents and could have distracted from the source. However, post-hoc tests including task repetition as a covariate and tests including only first judgments revealed the same pattern of results, rendering a strong bias towards social judgments, distracting from the sources due to task repetitions as unlikely. We can additionally show with eye-tracking that the source of the information is actively acknowledged during news exposure. We would also like to note that judging others based on visual appearance or minimal person-related information seems to be a natural tendency –we spontaneously form impressions about others and draw inferences about their character from minimal information (Bliss-Moreau et al., 2008; Foster, 2004; Todorov et al., 2007; Uhlmann et al., 2015). We therefore have no reason to assume that the results are due to the experimental situation. Indeed, in a short interview after the experiment (available from 29 participants), 27 expressed no doubt about the authenticity of the media reports. Taken together, our findings complement recent online studies on how true, misleading, or false information spreads and how news and its sources are evaluated (e.g. Brady et al., 2017; Pennycook & Rand, 2018, Vosoughi et al., 2018) by providing experimental insight into the precise neurocognitive mechanisms that underlie such behavior.

The current study was explicitly designed so that influences of the visual appearance of the faces were controlled for by careful counterbalancing. Facial trustworthiness can however influence person perception and memory (Lischke et al., 2018; Wendt et al., 2017; Weymar et al., 2019), and thus, it would be interesting to investigate how facial appearance-based information, such as trustworthiness may modulate the effects of emotional information and its credibility. First evidence suggests independence of emotional information and facial appearance (Eiserbeck & Abdel Rahman, 2020; Mattarozzi et al., 2014).

We conclude that the influence of source credibility on the effects of emotional contents of news headlines is remarkably weak. It is conceivable that source credibility did not qualify judgments because participants merely remembered the emotional content of the news but not the source (cf. Johnson et al., 1993; Yonelinas, 2002) or that they deliberately or unintentionally ignored the credibility of the source. This distinction cannot be made based on the current results and may be targeted in future studies. Future studies may identify the circumstances under which the influence of source credibility is strengthened. This may for example depend on how salient the source is and how clearly it is represented in memory and contextually available. Future research may further target emotion regulation (Gross, 2015; Maroney & Gross, 2014) and enhanced awareness about the consequences of potentially misleading information from sources of questionable credibility as a protection against biased social judgments.

## Author’s contribution

J. Baum and R. Abdel Rahman designed research and wrote the paper; J. Baum performed research and analyzed data.

## Acknowledgements

We thank Guido Kiecker for technical support and Cornelius Braun, Alexander Enge, Anna Faschinger, Kirsten Stark, and Martin Zielinski for assisting in stimulus preparation and/or data acquisition.

## Data Availability

Data and code is available to the reviewers and upon publication.

## Footnotes

2 Will be published upon peer-reviewed publication

3 LMMs yield the same pattern of results for ordinal dependent variables as CLMMs, please see SI-Table S2 for LMM results of ordinal dependent variables treated as continuous.

4 Will be published upon peer-reviewed publication

