## Supplemental Information for "Emotional news affects social judgments independent of perceived media credibility"

**Sample Size**

The following power analysis was preregistered on the OSF (Baum & Abdel Rahman, 2018[1]). The sample size was planned according to the counterbalancing of conditions, which requires a multiple of 6 participants, and based on expected coefficients for effects in electrophysiological brain responses (cf. Baum et al., 2018). We ran mixed model simulations (1000 for each coefficient, SIMR package in R, Green & MacLeod, 2016) to estimate the expected power for headline content effects and interactions with source credibility for a sample of 30 participants. Expected main effects of positive and negative headline content relative to neutral would be found with 100% power and respective interactions with source credibility of as small as 0.15μV would be found with over 90% power (**EPN**: grand mean of -0.4μV, effects of positive and negative headline content of 0.5μV each (power of 100%, 95% CI [99, 100]), interactions with source credibility of 0.15μV (power of 95.50%, 95% CI [94.02, 96.70] for negative vs. neutral headlines, power of 96.30%, 95% CI [94.94, 97.38] for positive vs. neutral headlines) were not expected but estimated in case they already occurred in the EPN; **LPP**: grand mean of 5μV, effects of headline content of 0.7μV (power of 100%, 95% CI [99, 100]), effect of source credibility of 0.3μV (possible but not expected), interactions with source credibility of 0.15μV (power of 95.50%, 95% CI [94.02, 96.70] for negative vs. neutral headlines, power of 96.30%, 95% CI [94.94, 97.38] for positive vs. neutral headlines)).

---

[1] https://osf.io/scbgq?view_only=1a818120b879472982eae0b75c251ad7

### Results

#### Effects of Emotional News on Information Processing and Social Judgments (Phase 2)

##### *Behavioral results.*

Table S1

*Means and confidence intervals for behavioural results of the social judgment task (on a 5-point scale of -2= very negative, 0 = neutral, 2 = very positive) using the SummarySEwithin function by Morey (2008) on single trial data.*

| Headline Source | Neutral Trusted | Neutral Distrusted | Negative Trusted | Negative Distrusted | Positive Trusted | Positive Distrusted |
|---|---|---|---|---|---|---|
| | | | **Social Judgement Decisions** | | | |
| *M* | 0.28 | 0.25 | -1.63 | -1.64 | 1.44 | 1.43 |
| 95%-CI | 0.25, 0.31 | 0.22, 0.28 | -1.66, -1.59 | -1.67, -1.6 | 1.41, 1.48 | 1.4, 1.47 |
| | | | **Social Judgment Latencies (ms)** | | | |
| M | 897.73 | 883.55 | 809.96 | 798.84 | 856.57 | 843.96 |
| 95%-CI | 883.32, 912.15 | 869.13, 897.97 | 796.75 823.18 | 786.43, 811.26 | 842.48, 870.67 | 830.98, 856.94 |

Table S2

*LMM summary statistics show effects of source credibility, negative and positive headline content and their interactions on social judgment decisions. Fixed effects were coded as repeated contrasts according to our hypotheses. The model is based on 14367 observations.*

| | **Social Judgment Decisions** | | | |
|---|---|---|---|---|
| *Coefficient* | *b (95%-CI)* | *SE* | *t* | *p* |
| Intercept (Grand Mean) | 0.02 (-0.08 – 0.12) | 0.05 | 0.45 | 0.656 |
| Source Credibility (Trusted vs. Distrusted) | 0.02 (-0.08 – 0.11) | 0.05 | 0.31 | 0.761 |
| Negative Headline Content (Neg vs. Neu) | -1.90 (-2.13 – -1.66) | 0.12 | -15.98 | <0.001 |
| Source Credibility * Negative Headline | -0.02 (-0.31 – 0.27) | 0.15 | -0.13 | 0.897 |
| Positive Headline Content (Pos vs. Neu) | 1.17 (0.94 – 1.40) | 0.12 | 10.01 | <0.001 |
| Source Credibility * Positive Headline | -0.02 (-0.24 – 0.19) | 0.11 | -0.22 | 0.826 |
| Model Formula | Decision ~ Headline Content * Source Credibility + (S+Neg+S*Neg+Pos+S*Pos \| subject) + (S+Neg+S*Neg+Pos+S*Pos \| face) | | | |

*Note.* Abbreviations for slopes in the random effects terms: S=Source Credibility, Neg=Negative Headline Content, Pos=Positive Headline Content. Face stands for face stimulus.

Table S3

*Mixed model summary statistics for social judgment decisions and latencies including the repetition of task as a covariate. Each model is based on 14367 observations.*

| | **Social Judgment Decisions** | | | | **Social Judgment Latencies [-1000/latency(ms)]** | | | |
|---|---|---|---|---|---|---|---|---|
| *Coefficient* | *b* | *SE* | *z* | *p* | *b (95%-CI)* | *SE* | *t* | *p* |
| Intercept (Grand Mean) | | | | | -1.08 (-1.14 – -1.03) | 0.03 | -36.35 | <0.001 |
| Repetition | -0.00 | 0.00 | -0.03 | 0.975 | -0.02 (-0.02 – -0.02) | 0.00 | -51.14 | <0.001 |
| Source Credibility (Trusted vs. Distrusted) | 0.02 | 0.40 | 0.06 | 0.955 | 0.02 (-0.02 – 0.05) | 0.02 | 0.99 | 0.324 |
| Source Credibility * Rep | -0.00 | 0.00 | -0.38 | 0.702 | -0.00 (-0.00 – 0.00) | 0.00 | -0.58 | 0.560 |
| Negative Headline Content (Neg vs. Neu) | -8.13 | 0.75 | -10.80 | <0.001 | -0.17 (-0.23 – -0.12) | 0.03 | -5.92 | <0.001 |
| Source Credibility * Negative Headline Content | -0.39 | 1.13 | -0.35 | 0.726 | -0.02 (-0.09 – 0.06) | 0.04 | -0.46 | 0.645 |
| Negative Headline Content * Rep | 0.00 | 0.00 | -0.96 | 0.337 | 0.00 (0.00 – 0.01) | 0.00 | 4.30 | <0.001 |
| Negative Headline Content * Source Credibility * Rep | 0.01 | 0.02 | 0.58 | 0.565 | -0.00 (-0.00 – 0.00) | 0.00 | -0.11 | 0.915 |
| Positive Headline Content (Pos vs. Neu) | 4.56 | 0.62 | 7.38 | <0.001 | -0.08 (-0.12 – -0.03) | 0.02 | -3.43 | 0.001 |
| Source Credibility * Positive Headline Content | -0.03 | 0.49 | -0.06 | 0.953 | -0.03 (-0.11 – 0.05) | 0.04 | -0.76 | 0.448 |
| Positive Headline Content * Rep | 0.04 | 0.00 | 4.38 | <0.001 | 0.00 (-0.00 – 0.00) | 0.00 | 1.63 | 0.103 |
| Positive Headline Content * Source Credibility * Rep | -0.00 | 0.02 | -0.15 | 0.878 | 0.00 (-0.00 – 0.01) | 0.00 | 0.58 | 0.564 |
| Model Formula | Decision ~ Headline Content * Source Credibility * Rep + (S+Neg+S*Neg+Pos+S*Pos \|\| subject) + (S+Neg+S*Neg+Pos+S*Pos \|\| face) | | | | Latency ~ Headline Content * Source Credibility * Rep + (S+Neg+S*Neg+Pos+S*Pos \|\| subject) + (S+Neg+S*Neg+Pos+S*Pos \|\| face) | | | |

*Note.* Double bars in random effects terms set correlation parameters to zero. Abbreviations for slopes in the random effects terms: S=Source Credibility, Neg=Negative Headline Content, Pos=Positive Headline Content. Face stands for face stimulus.
CLMM threshold coefficients for social judgment decisions: -2 | -1: *b*=-3.9, *SE*=.28, *z*=-50.0; -1 | 0: *b*=-1.76, *SE*=.07, *z*=-26.7; 0 | 1 *b*=2.0, *SE*=.06, *z*=31.8; 1 | 2: *b*=5.1, *SE*=.08, *z*=64.5

Table S4

*Mixed model summary statistics for for social judgment decisions and latencies testing only the first repetition of the task based on 713 observations.*

| | **Social Judgment Decisions** | | | | **Social Judgment Latencies [-1000/latency(ms)]** | | | |
|---|---|---|---|---|---|---|---|---|
| *Coefficient* | *b* | *SE* | *z* | *p* | *b (95%-CI)* | *SE* | *t* | *p* |
| Intercept (Grand Mean) | | | | | -0.96 (-1.02 – -0.89) | 0.03 | -28.48 | <0.001 |
| Source Credibility (Trusted vs. Distrusted) | -0.10 | 0.21 | -0.49 | 0.627 | 0.02 (-0.02 – 0.06) | 0.02 | 1.02 | 0.324 |
| Negative Headline Content (Neg vs. Neu) | -5.39 | 0.53 | -10.10 | <0.001 | -0.19 (-0.26 – -0.12) | 0.04 | -5.50 | <0.001 |
| Source Credibility * Negative Headline Content | -0.15 | 0.43 | -0.34 | 0.735 | -0.05 (-0.14 – 0.04) | 0.05 | -1.07 | 0.297 |
| Positive Headline Content (Pos vs. Neu) | 3.01 | 0.42 | 7.15 | <0.001 | -0.10 (-0.14 – -0.06) | 0.02 | -5.20 | <0.001 |
| Source Credibility * Positive Headline Content | 0.31 | 0.44 | 0.69 | 0.491 | -0.02 (-0.11 – 0.08) | 0.05 | -0.34 | 0.734 |
| Model Formula | Decision ~ Headline Content * Source Credibility (S+Neg+Pos+S*Pos \|\| subject) + (S+Neg+S*Neg+Pos \|\| face) | | | | Latency ~ Headline Content * Source Credibility + (S+Neg+S*Neg \|\| subject) + ( S+Neg+S*Neg+ S*Pos \|\| face) | | | |

*Note.* Double bars in random effects terms set correlation parameters to zero. Abbreviations for slopes in the random effects terms: S=Source Credibility, Neg=Negative Headline Content, Pos=Positive Headline Content. Face stands for face stimulus.

CLMM threshold coefficients for social judgment decisions: -2 | -1: *b*=-3.1, *SE*=.22, *z*=-14.2; -1 | 0: *b*=-1.24, *SE*=.16, *z*=-7.9; 0 | 1 *b*=1.36, *SE*=.15, *z*=9.2; 1 | 2: *b*=3.6, *SE*=.20, *z*=17.7

***Event-related brain potentials.***

Table S5

*Means and confidence intervals for the EPN and LPP amplitudes in microvolts averaged over respective ROIs and time windows using the SummarySEwithin function by Morey (2008) on single trial data*

| Headline Source | Neutral Trusted | Neutral Distrusted | Negative Trusted | Negative Distrusted | Positive Trusted | Positive Distrusted |
|---|---|---|---|---|---|---|
| | | | **EPN** | | | |
| *M* | 2.42 | 2.66 | 2.36 | 2.15 | 2.4 | 2.46 |
| 95%-CI | 2.23 – 2.60 | 2.47 – 2.85 | 2.17 – 2.54 | 1.96– 2.34 | 2.21– 2.59 | 2.26– 2.65 |
| | | | **LPP** | | | |
| *M* | 4.07 | 4.17 | 5.38 | 5.09 | 4.66 | 4.54 |
| 95%-CI | 3.87 – 4.27 | 3.97 – 4.37 | 5.17 – 5.59 | 4.88 – 5.29 | 4.45 – 4.86 | 4.34 – 4.73 |

Table S6

*LMM results for the LPP including judgment latencies (RT) as covariate based on 1451 observations. RT were centered and multiplied by 0.01 to get similar scaled predictors.*

| | | | **LPP** | | |
|---|---|---|---|---|---|
| *Coefficient* | *b* | *SE* | *95%-CI* | *t* | *p* |
| Intercept (Grand Mean) | 4.62 | 0.38 | 3.88 – 5.37 | 12.16 | <0.001 |
| RT | -0.17 | 0.01 | -0.19 – -0.14 | -13.52 | <0.001 |
| Source Credibility (Trusted vs. Distrusted) | 0.13 | 0.10 | -0.06 – 0.32 | 1.31 | 0.206 |
| Source Credibility * RT | 0.01 | 0.02 | -0.03 – 0.06 | 0.60 | 0.551 |
| Negative Headline Content (Neg vs. Neu) | 0.97 | 0.16 | 0.66 – 1.28 | 6.22 | <0.001 |
| Source Credibility * Negative Headline Content | 0.38 | 0.20 | -0.01 – 0.77 | 1.90 | 0.065 |
| Negative Headline Content * RT | -0.14 | 0.03 | -0.20 – -0.08 | -4.72 | <0.001 |
| Negative Headline Content * Source Credibility * RT | 0.06 | 0.06 | -0.06 – 0.17 | 1.00 | 0.318 |
| Positive Headline Content (Pos vs. Neu) | 0.45 | 0.13 | 0.19 – 0.71 | 3.43 | 0.002 |
| Source Credibility * Positive Headline Content | 0.21 | 0.25 | -0.28 – 0.69 | 0.84 | 0.409 |
| Positive Headline Content * RT | -0.04 | 0.03 | -0.09 – 0.02 | -1.32 | 0.187 |
| Positive Headline Content * Source Credibility * RT | 0.04 | 0.06 | -0.07 – 0.15 | 0.67 | 0.500 |
| Model Formula | LPP ~ Headline Content * Source Credibility * RTcentered<br>+ (S+Neg+S*Neg+Pos+S*Pos \|\| subject)<br>+ (S+Neg +Pos+S*Pos \|\| face) | | | | |

*Note.* Double bars in random effects terms set correlation parameters to zero. Abbreviations for slopes in the random effects terms: S=Source Credibility, Neg=Negative Headline Content, Pos=Positive Headline Content. Face stands for face stimulus.

#### *Visual perception in the P100 and N170 (additional post-hoc analyses)*

Additionally to the EPN and LPP, we explored possible effects during earlier stages of visual face perception in the P100 (Tables S12 and S13) and N170 (Table S14 and S15). Compared to neutral headlines, there were no effects of negative or positive headlines in the P100 or N170. However, the effect of negative compared to neutral headlines in the N170 interacted with source credibility, due to an enhanced N170 for distrusted sources, and an absent effect for trusted sources.

Table S7

*Means and confidence intervals for the P100 and N170 amplitudes in microvolts averaged over respective ROIs and time windows using the SummarySEwithin function by Morey (2008) on single trial data.*

| Headline<br>Source | Neutral<br>Trusted | Neutral<br>Distrusted | Negative<br>Trusted | Negative<br>Distrusted | Positive<br>Trusted | Positive<br>Distrusted |
|---|---|---|---|---|---|---|
| | | | **P100** | | | |
| *M* | 3.10 | 3.16 | 3.29 | 3.35 | 3.22 | 3.29 |
| 95%-CI | 2.90 – 3.31 | 2.95 – 3.36 | 3.09 – 3.50 | 3.15 – 3.56 | 3.01 – 3.43 | 3.09 – 3.50 |
| | | | **N170** | | | |
| *M* | -2.04 | -1.82 | -1.95 | -2.14 | -2.09 | -1.94 |
| 95%-CI | -2.24 – -1.84 | -2.02 – -1.63 | -2.15 – -1.75 | -2.34 – -1.95 | -2.29 – -1.89 | -2.14 – -1.73 |

Table S8

*LMM summary statistics show effects of source credibility, negative and positive headline content and their interactions on P1 and N170 as dependent variables in the social judgment task. Effects on the ROI and time range of the P1 and N170 amplitudes were estimated in separate LMMs and fixed effects were coded as repeated contrasts. Each model is based on 14151 observations.*

| | **P1** | | | | **N170** | | | |
|---|---|---|---|---|---|---|---|---|
| *Coefficient* | *b (95%-CI)* | *SE* | *t* | *p* | *b (95%-CI)* | *SE* | *t* | *p* |
| Intercept (Grand Mean) | 3.24 (2.17 – 4.31) | 0.54 | 5.95 | <0.001 | -1.99 (-3.09 – -0.89) | 0.56 | -3.55 | 0.001 |
| Source Credibility (Trusted vs. Distrusted) | -0.06 (-0.23 – 0.12) | 0.09 | -0.66 | 0.514 | -0.06 (-0.26 – 0.15) | 0.10 | -0.54 | 0.592 |
| Negative Headline Content (Neg vs. Neu) | 0.20 (-0.06 – 0.45) | 0.13 | 1.53 | 0.138 | -0.11 (-0.30 – 0.07) | 0.09 | -1.22 | 0.231 |
| Source Credibility * Negative Headline Content | -0.03 (-0.42 – 0.37) | 0.20 | -0.13 | 0.900 | 0.40 (0.03 – 0.78) | 0.19 | 2.10 | 0.041 |
| Positive Headline Content (Pos vs. Neu) | 0.13 (-0.13 – 0.39) | 0.13 | 1.00 | 0.326 | -0.08 (-0.31 – 0.15) | 0.12 | -0.66 | 0.515 |
| Source Credibility * Positive Headline Content | -0.06 (-0.58 – 0.47) | 0.27 | -0.21 | 0.838 | 0.04 (-0.41 – 0.48) | 0.23 | 0.15 | 0.878 |
| Model Formula | P100 ~ Headline Content * Source Credibility + (Neg+S*Neg+Pos+S*Pos \|\| subject) + (S+Neg+Pos \|\| face) | | | | N170 ~ Headline Content * Source Credibility + (S+Neg+S*Neg+Pos \|\| subject) + (S+Pos+S*Pos \|\| face) | | | |

*Note.* Double bars in random effects terms set correlation parameters to zero. Abbreviations for slopes in the random effects terms: S=Source Credibility, Neg=Negative Headline Content, Pos=Positive Headline Content. Face stands for face stimulus.

Table S9

*Negative and positive headline content effects on P1 and N170 separately within each source credibility condition computed from the models in Table 7. Each model is based on 14151 observations.*

| | **P1** | | | | **N170** | | | |
|---|---|---|---|---|---|---|---|---|
| *Contrast* | *b* | *SE* | *t* | *p* | *b* | *SE* | *t* | *p* |
| Trusted: Neg vs. Neu | 0.19 | 0.17 | 1.12 | 0.532 | 0.09 | 0.14 | 0.65 | 0.715 |
| Distrusted: Neg vs. Neu | 0.21 | 0.17 | 1.28 | 0.532 | -0.32 | 0.14 | -2.35 | 0.084 |
| Trusted: Pos vs. Neu | 0.10 | 0.19 | 0.55 | 0.584 | -0.06 | 0.16 | -0.37 | 0.715 |
| Distrusted: Pos vs. Neu | 0.16 | 0.19 | 0.85 | 0.535 | -0.10 | 0.16 | -0.58 | 0.715 |

Fig. S1: **Phase 2** Main task: Social judgment; N170 results.

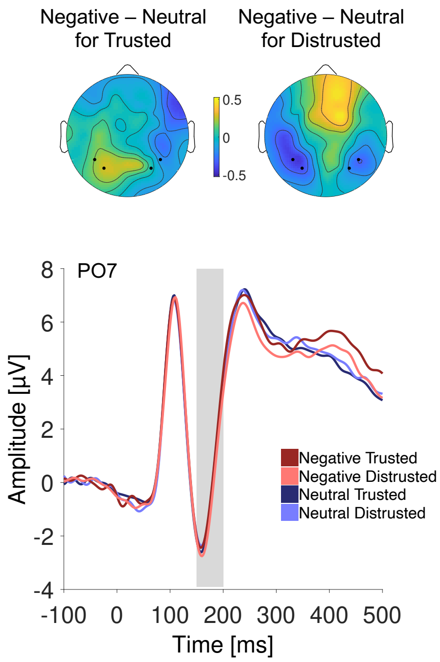


Fig. S1. In **Phase 2** the EEG was acquired while a social judgment task was employed as the main task to investigate the effects of emotional news. ERP time point zero was the face onset on the screen. Results show an N170 (150–200ms) effect for persons related to negative headline content (compared to neutral headline content) from distrusted sources. Grand average ERPs are shown at electrode site PO7, and scalp distributions show the effect as difference between conditions in the time window shaded in grey.

**News Exposure and Manipulation Checks (Phase 1)**

Table S10

*Means and confidence intervals for the person likability ratings pre- and post-exposure on a 5-Point Scale (-2= very unlikable, 0 = neutral, 2 = very likable) using the SummarySEwithin function by Morey (2008) on single trial data.*

| Headline Source | Neutral Trusted | Neutral Distrusted | Negative Trusted | Negative Distrusted | Positive Trusted | Positive Distrusted |
|---|---|---|---|---|---|---|
| Before Exposure | | | | | | |
| *M* | -0.12 | 0.18 | -0.1 | -0.05 | -0.02 | -0.02 |
| 95%-CI | -0.28, 0.05 | 0.01, 0.36 | -0.26, 0.06 | -0.24, 0.14 | -0.19, 0.15 | -0.19, 0.16 |
| After Exposure | | | | | | |
| M | 0.29 | 0.22 | -1.28 | -1.24 | 1.12 | 0.95 |
| 95%-CI | 0.13, 0.45 | 0.06, 0.39 | -1.47, -1.1 | -1.42, -1.06 | 0.96, 1.29 | 0.75, 1.15 |

Table S11

*CLMM summary statistics for the person likability ratings pre and post-exposure based on 1440 observations. The predictors headline content and source credibility were nested in pre- and post-exposure (coded as before vs. after).*

| | **Likability Rating Pre and Post News Exposure** | | | |
|---|---|---|---|---|
| *Coefficient* | *b* | *SE* | *z* | *p* |
| Pre vs. Post | -0.02 | 0.10 | -0.17 | 0.863 |
| Pre: Source Credibility (Trusted vs. Distrusted) | -0.22 | 0.15 | -1.43 | 0.153 |
| Pre: Negative Headline Content (Neg vs. Neu) | -0.23 | 0.29 | -0.80 | 0.427 |
| Pre: Source Credibility * Negative Headline Content | 0.52 | 0.33 | 1.55 | 0.122 |
| Pre: Positive Headline Content (Pos vs. Neu) | -0.08 | 0.19 | -0.42 | 0.672 |
| Pre: Source Credibility * Positive Headline Content | 0.56 | 0.33 | 1.69 | 0.091 |
| Post: Source Credibility (Trusted vs. Distrusted) | 0.04 | 0.16 | 0.22 | 0.823 |
| Post: Negative Headline Content (Neg vs. Neu) | -3.28 | 0.31 | -10.57 | <0.001 |
| Post: Source Credibility * Negative Headline Content | -0.27 | 0.35 | -0.78 | 0.435 |
| Post: Positive Headline Content (Pos vs. Neu) | 1.79 | 0.20 | 8.98 | <0.001 |
| Post: Source Credibility * Positive Headline Content | 0.22 | 0.34 | 0.65 | 0.516 |
| Model Formula | Rating ~ Phase/(Headline Content * Source Credibility)<br>+ (S+Neg +Pos \|\| subject)<br>+ (S+Neg +Pos \|\| face) | | | |

*Note.* Double bars in random effects terms set correlation parameters to zero. Abbreviations for slopes in the random effects terms: S=Source Credibility, Neg=Negative Headline Content, Pos=Positive Headline Content. Face stands for face stimulus.

CLMM threshold coefficients: -2 | -1: *b*=-2.75, *SE*=.11, *z*=-25.1; -1 | 0: *b*=-0.73, *SE*=.07, *z*=-11.1; 0 | 1 *b*=0.74, *SE*=.06, *z*=11.4; 1 | 2: *b*=2.97, *SE*=.11, *z*=-27.1

Table S12

*Negative and positive headline content effects on person likability ratings post-exposure separately within each source credibility condition computed from the model in Table S10. Each model is based on 1440 observations.*

| | **Likability Rating Post News Exposure** | | | |
|---|---|---|---|---|
| *Contrast* | *b* | *SE* | *z* | *p* |
| Trusted: Neg vs. Neu | -3.88 | 0.61 | -6.39 | <0.001 |
| Distrusted: Neg vs. Neu | -3.52 | 0.60 | -5.91 | <0.001 |
| Trusted: Pos vs. Neu | 2.16 | 0.34 | 6.37 | <0.001 |
| Distrusted: Pos vs. Neu | 1.92 | 0.34 | 5.99 | <0.001 |

Table S13

*Means and confidence intervals for recognition accuracy in the post-exposure recognition test using the SummarySEwithin function by Morey (2008) on single trial data.*

| Headline Source | Neutral Trusted | Neutral Distrusted | Negative Trusted | Negative Distrusted | Positive Trusted | Positive Distrusted |
|---|---|---|---|---|---|---|
| *M* | 0.98 | 0.98 | 0.96 | 0.94 | 1 | 0.99 |
| 95%-CI | 0.95 – 1 | 0.94 – 1 | 0.92– 1 | 0.90 – 0.99 | 0.99 – 1 | 0.97 – 1 |

Table S14

*GLMM summary statistics for recognition accuracy in the post-exposure recognition test based on 716 observations.*

| | Correctly recognized | | | |
|---|---|---|---|---|
| *Coefficient* | b | *SE* | *z*-value | *p* |
| Intercept (Grand Mean) | 6.37 | 71.45 | 0.09 | 0.929 |
| Source Credibility (Trusted vs. Distrusted) | 4.10 | 142.89 | 0.03 | 0.977 |
| Negative Headline Content (Neg vs. Neu) | -0.75 | 0.53 | -1.33 | 0.184 |
| Source Credibility * Negative Headline Content | 0.38 | 1.14 | 0.34 | 0.737 |
| Positive Headline Content (Pos vs. Neu) | 7.11 | 215.34 | 0.03 | 0.974 |
| Source Credibility * Positive Headline Content | 11.88 | 428.68 | 0.03 | 0.978 |
| Model Formula | CorrRec ~ Headline Content * Source Credibility + (1\| subject) + (1\| face) | | | |

*Note.* GLMM (binomial data) was used. Abbreviations for slopes in the random effects terms: S=Source Credibility, Neg=Negative Headline Content, Pos=Positive Headline Content. Face stands for face stimulus.

***Eye Tracking Experiment.***

Websites were presented in a 10% reduced size (1152 × 922 px) on a grey background to prevent excess data loss at the edges of the screen, which would have been possible if the stimuli were presented full screen (1024 × 1280 px). The eye tracker (SMI Red, SensoMotoric Instruments, Inc., Boston, MA) was calibrated before the experiment and if necessary, an additional offset correction was performed offline during the presentation of a fixation cross in the beginning of the news exposure. Fixations were detected using default event detection algorithms of SMI's BeGaze software 3.5 (min. duration 80ms, max. dispersion 100px). We measured fixation durations and frequencies in the areas where the media source logos were presented. The area of interest for the media source logo was the same size for all logos (641px × 266 px), being large enough to cover the slightly varying spatial extensions of the different logos and was placed on every stimulus individually so that it did not include possible other distractors (e.g. salient parts of the blurred website). Fixation durations and frequencies within the logo regions were accumulated by face stimulus across the five news report presentations during the exposure phase within participants, and averaged across face stimuli. Filler trials were excluded.

**News Media Source Checks (Phase 3)**

Table S15

*Means and confidence intervals for source trustworthiness and likability ratings on 5-Point Scales (-2= untrustworthy / unlikable, 0 = neutral, 2 = trustworthy/ likable) using the SummarySEwithin function by Morey (2008) on single trial data.*

| Rating | Trustworthiness Rating | | Likability Rating | |
|---|---|---|---|---|
| Source | Trusted | Distrusted | Trusted | Distrusted |
| *M* | 1.41 | -1.61 | 1.13 | -1.43 |
| 95%-CI | 1.36 – 1.46 | -1.66 – -1.56 | 0.94 – 1.32 | -1.63 – -1.22 |

Table S16

*CLMM summary statistics for source trustworthiness and likability ratings based on 2400 (trustworthiness) and 240 (likability) observations.*

| | Trustworthiness Rating | | | | Likability Rating | | | |
|---|---|---|---|---|---|---|---|---|
| *Coefficient* | *b* | *SE* | *z* | *p* | *b* | *SE* | *z* | *p* |
| Source Credibility (Trusted vs. Distrusted) | 10.63 | 0.96 | 11.02 | <0.001 | 6.47 | 0.93 | 6.99 | <0.001 |
| Model Formula | Rating ~ Source Credibility + (S \| subject) + (1 \| media stimulus) | | | | Rating ~ Source Credibility + (S \| subject) + (1 \| media stimulus) | | | |

*Note.* Abbreviation used in the random effects terms: S=Source Credibility.
CLMM threshold coefficients for trustworthiness: -2 | -1: *b*=-3.8, *SE*=0.50, *z*=-7.56; -1 | 0: *b*=-0.7, *SE*=0.05, *z*=-1.4; 0 | 1 *b*=1.1, *SE*=0.50, *z*=2.2; 1 | 2: *b*=5.3, *SE*=0.51, *z*=10.34
CLMM threshold coefficients for likability: -2 | -1: *b*=-2.6, *SE*=0.48, *z*=-5.40; -1 | 0: *b*=-0.8, *SE*=0.46, *z*=-1.7; 0 | 1 *b*=1.4, *SE*=0.47, *z*=2.9; 1 | 2: *b*=3.8, *SE*=0.53, *z*=7.2

***ERPs evoked by News Media Logos during Source Trustworthiness Ratings***

To additionally investigate the processing of news media sources, we recorded the EEG while logos were presented and participants rated how trustworthy they consider each source. While differences in visual features of the logos (e.g. color) may confound these effects, the effects remain significant when accounting for the average responses evoked by individual logos in the LMM. The EPN was more negative for distrusted than trusted sources (see Table S17). There was no effect in the LPP (see Table S17).

Table S17

*Means and confidence intervals of the EPN (EPN ROI and time window) and the LPP (LPP ROI and time window) evoked by news media logos in microvolts using the SummarySEwithin function by Morey (2008) on single trial data.*

| Rating | EPN | | LPP | |
|---|---|---|---|---|
| Source | Trusted | Distrusted | Trusted | Distrusted |
| *M* | 0.57 | -0.53 | 4.35 | 4.54 |
| 95%-CI | 0.22 – 0.91 | -0.88 – -0.19 | 4 – 4.71 | 4.17 – 4.92 |

Table S18

*LMM summary statistics for the EPN and the LPP evoked by news media source logos based on 2358 observations.*

| | EPN | | | | | LPP | | | | |
|---|---|---|---|---|---|---|---|---|---|---|
| *Coefficient* | *b* | *95%-CI* | *SE* | *t* | *p* | *b* | *95%-CI* | *SE* | *t* | *p* |
| Intercept (Grand Mean) | 0.02 | -0.96 – 0.99 | 0.50 | 0.03 | 0.974 | 4.44 | 3.87 – 5.02 | 0.29 | 15.14 | <0.001 |
| Source Credibility (Trusted vs. Distrusted) | 1.11 | 0.22 – 2.01 | 0.46 | 2.43 | 0.037 | -0.18 | -0.68 – 0.31 | 0.25 | -0.73 | 0.480 |
| Model Formula | | EPN ~ Source Credibility + (S\|subject) + (1\|media stimulus) | | | | | LPP ~ Source Credibility + (S\|subject) + (1\|media stimulus) | | | |

*Note.* Abbreviation used in the random effects terms: S=Source Credibility.

Fig. S2: **Phase 3** EPN evoked by logos of news media sources during trustworthiness ratings.

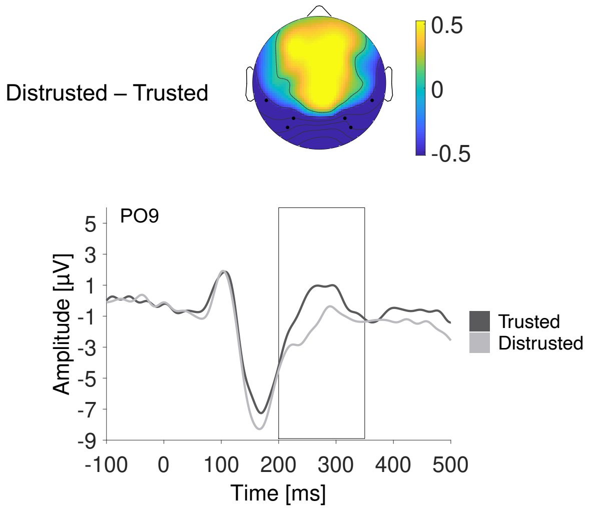


Fig. S2. In **Phase 3** the logos of news media sources were presented and the EEG was acquired while participants rated the sources' trustworthiness. ERP time point zero was the logo onset on the screen. Results show an EPN (200–350ms) effect for logos from distrusted sources compared to trusted sources. Grand average ERPs are shown at electrode site PO9, and scalp distributions show the effect as difference between conditions in the time window within the box.

### Materials

Twenty-four unfamiliar faces with neutral facial expression were taken from multiple databases, for example Ebner, Riediger, & Lindenberger (2010) and Langner et al. (2010).

Headlines were pre-rated regarding valence and arousal on 7-point scales (participants were $N = 17$; 7 females, mean age 26 ($SD = 2.28$), all students). Negative and positive headlines significantly differed from neutral in terms of valence, $t(16) = -14.59$, $p < .001$, and $t(16) = 15.09$, $p < .001$ respectively, as well as in terms of arousal, $t(16) = 10.31$, $p < .001$, and $t(16) = 11.25$, $p < .001$ respectively. Positive and negative headlines were equally arousing, $t(16) = .07$, $p = .94$.

Table S19

*Pre-rating means and 95% confidence intervals for headlines.*

| | Negative | Positive | Neutral |
|---|---|---|---|
| Valence *M*, 95% CI | 1.99 [1.74, 2.25] | 5.96 [5.70, 6.23] | 4.02 [3.95, 4.09] |
| Arousal *M*, 95% CI | 4.33 [3.80, 4.87] | 4.31 [3.75, 4.88] | 1.67 [1.31, 2.02] |

Table S20

*Headlines used in the exposure phase with pre-ratings of valence and arousal on a 7-point scale from 1 (very negative; not arousing) to 7 (very positive, very arousing). German originals with English translations. The headlines had two parts as that was required by the layouts of the news websites and referred to the person depicted.*

| **Negative headlines** | Valence *M*, (*SD*) | Arousal *M*, (*SD*) |
|---|---|---|
| Steuergelder veruntreut - Diese Abgeordnete verschafft ihrem Ehemann Beamtengehalt<br>*Taxpayers' money embezzled - This MP provides her husband with civil servant's salary* | 2.18<br>(1.19) | 3.71<br>(1.45) |
| Berlin Neukölln - Dieser Casinobetreiber hat Jugendlichen zum Kokainschmuggel gezwungen<br>*Berlin Neukölln - This casino owner forced teenager to smuggle cocaine* | 1.82<br>(.809) | 4.59<br>(1.28) |
| Abzocke - Sie gab sich als Putzhilfe aus, um Senioren zu bestehlen<br>*Rip-off - She pretended to be a cleaner to steal from seniors* | 2.47<br>(.624) | 3.65<br>(1.54) |
| Tatort Kneipe - Er mischte seinen Verabredungen K.O.-Tropfen ins Getränk<br>*Crime scene "Pub" - He spiked the drinks of his dates* | 1.71<br>(.686) | 4.59<br>(1.70) |
| Position ausgenutzt - Diese Frau tyrannisierte ihren Lehrling gezielt über mehrere Monate<br>*Position exploited - This woman deliberately tormented her apprentice over several months* | 2.18<br>(.809) | 4.12<br>(1.54) |
| Kinder in Gefahr - Dieser Mann hat harte Drogen auf dem Schulhof verkauft<br>*Children in danger - This man sold hard drugs in the schoolyard* | 2.00<br>(.707) | 4.35<br>(1.58) |
| Organisierte Kriminalität - Ich beklaue mit meiner Familie Touristen im großen Stil<br>*Organized crime – Me and my family are robbing tourists on a grand scale* | 2.12<br>(.993) | 4.65<br>(1.41) |
| Hetze in Frankfurt - Ich habe Israelis schon immer gehasst<br>*Baiting in Frankfurt - I have always hated Israelis* | 1.47<br>(.514) | 5.00<br>(1.23) |
| Mean values of all negative headlines | 1.99<br>(.791) | 4.33<br>(1.46) |

| Positive headlines | Valence *M*, (*SD*) | Arousal *M*, (*SD*) |
|---|---|---|
| Erfolg der Wissenschaft - Diese Forscherin schenkt vielen Erblindeten das Augenlicht zurück<br>*Success of Science - This researcher gives back eyesight to many blind people* | 6.35 (.606) | 4.59 (1.42) |
| Fliegerische Meisterleistung - Dieser Pilot rettet Leben seiner Passagiere dank geglückter Notlandung<br>*Flying masterpiece - This pilot saves the lives of his passengers thanks to a successful emergency landing* | 6.12 (.781) | 4.53 (1.23) |
| Zivilcourage - Sie hat Mut bewiesen und einen Reisenden am Bahnsteig wiederbelebt<br>*Hero – She revived stranger on the train platform* | 6.12 (.697) | 4.35 (1.46) |
| Heldentat im Mittelmeer - Er rettete eine Familie vor dem Ertrinken<br>*Heroic act in the Mediterranean - He saved a family from drowning* | 6.06 (.827) | 4.41 (1.66) |
| Herz für Berlin - Diese Frau bewahrte Obdachlosen mit ihrem Wohnmobil vor dem Erfrieren<br>*Love for Berlin - This woman saved homeless by opening up her campervan* | 6.29 (.686) | 4.18 (1.38) |
| Passanten in Sicherheit - Dieser Mann stellte sich dem Angreifer in den Weg<br>*Bystanders safed - This man stood in the way of the attacker* | 5.35 (.786) | 4.29 (1.57) |
| Kinderschutz - Ich leite eine Hilfsorganisation zur Rettung von Kindersoldaten<br>*Child protection - I run an aid organization for the rescue of child soldiers* | 5.53 (.624) | 4.18 (1.33) |
| Freundschaft in Gaza - Ich habe Frieden mit meinen Nachbarn geschlossen<br>*Friendship in Gaza - I have made peace with my neighbors* | 5.88 (.928) | 4.00 (1.62) |
| Mean values of all positive headlines | 5.96 (.742) | 4.32 (1.46) |

| **Neutral headlines** | Valence *M*, (*SD*) | Arousal *M*, (*SD*) |
|---|---|---|
| Internetsicherheit erweitert - Diese Angestellte erklärt die neuen Leitlinien<br>*Internet safety enhanced - This employee explains the new guidelines* | 4.24<br>(.437) | 2.00<br>(1.17) |
| Abitur - Dieser Lehrer wurde als Vertreter in den Reformausschuss gewählt<br>*Graduation reform - This teacher was elected as a representative in the committee* | 3.94<br>(.243) | 1.35<br>(.862) |
| Personalwechsel - Sie hat die Leitung der Landesgartenschau für ein Jahr übernommen<br>*Change of CEO - She took the management of the state garden exhibition for one year* | 4.12<br>(.485) | 1.41<br>(.795) |
| Startup-Szene - Er investiert in smarte Haushaltsgeräte<br>*Startup scene - He invests in smart household appliances* | 4.00<br>(.612) | 1.88<br>(1.17) |
| Talbahn Kärnten - Dieser Mann war mit neuen Auftraggebern der Zugstrecke im Gespräch<br>*Valley railway Kärnten - This man was in conversation with the new contractors* | 4.06<br>(.556) | 1.41<br>(1.00) |
| Wintersport - Diese Frau ist die Newcomerin unter den Langläuferinnen<br>*Winter sports - This woman is the newcomer among cross-country skiers* | 4.24<br>(.562) | 1.94<br>(1.35) |
| Verkehr und Mobilität - Ich habe den neuen ADAC-Ratgeber für Autofahrer getestet<br>*Traffic and mobility - I tested the new automobile club's car guide* | 3.88<br>(.485) | 1.47<br>(.874) |
| Konsortium tagt - Ich habe zähe Verhandlungen bereits erwartet.<br>*Consortium meets - I was expecting tough negotiations* | 3.71<br>(.772) | 1.88<br>(1.05) |
| Mean values of all neutral headlines | 4.02<br>(.519) | 1.67<br>(1.03) |
